# Potent human neutralizing antibodies elicited by SARS-CoV-2 infection

**DOI:** 10.1101/2020.03.21.990770

**Authors:** Bin Ju, Qi Zhang, Xiangyang Ge, Ruoke Wang, Jiazhen Yu, Sisi Shan, Bing Zhou, Shuo Song, Xian Tang, Jinfang Yu, Jiwan Ge, Jun Lan, Jing Yuan, Haiyan Wang, Juanjuan Zhao, Shuye Zhang, Youchun Wang, Xuanling Shi, Lei Liu, Xinquan Wang, Zheng Zhang, Linqi Zhang

## Abstract

The pandemic caused by emerging coronavirus SARS-CoV-2 presents a serious global public health emergency in urgent need of prophylactic and therapeutic interventions. SARS-CoV-2 cellular entry depends on binding between the viral Spike protein receptor-binding domain (RBD) and the angiotensin converting enzyme 2 (ACE2) target cell receptor. Here, we report on the isolation and characterization of 206 RBD-specific monoclonal antibodies (mAbs) derived from single B cells of eight SARS-CoV-2 infected individuals. These mAbs come from diverse families of antibody heavy and light chains without apparent enrichment for particular families in the repertoire. In samples from one patient selected for further analyses, we found coexistence of germline and germline divergent clones. Both clone types demonstrated impressive binding and neutralizing activity against pseudovirus and live SARS-CoV-2. However, the antibody neutralizing potency is determined by competition with ACE2 receptor for RBD binding. Surprisingly, none of the SARS-CoV-2 antibodies nor the infected plasma cross-reacted with RBDs from either SARS-CoV or MERS-CoV although substantial plasma cross-reactivity to the trimeric Spike proteins from SARS-CoV and MERS-CoV was found. These results suggest that antibody response to RBDs is viral species-specific while that cross-recognition target regions outside the RBD. The specificity and neutralizing characteristics of this plasma cross-reactivity requires further investigation. Nevertheless, the diverse and potent neutralizing antibodies identified here are promising candidates for prophylactic and therapeutic SARS-CoV-2 interventions.

## Introduction

The source of the recent Coronavirus Disease 2019 (COVID-19) outbreak in Wuhan, China is a novel pathogenic coronavirus, SARS-CoV-2 ^1-4^. Its unique pathogenesis and rapid international transmission poses a serious global health emergency ^5-9^. SARS-CoV-2 belongs to the betacoronavirus family and shares substantial genetic and functional similarity with other pathogenic human betacoronaviruses, including Severe Acute Respiratory Syndrome Coronavirus (SARS-CoV) and Middle East Respiratory Syndrome Coronavirus (MERS-CoV) ^2-4,10,11^. The virus is believed to have originated in bats, although the source and animal reservoirs of SARS-CoV-2 remain uncertain ^2-4,10-12^. No SARS-CoV-2-specific treatments or vaccine are currently available but several antiviral drugs including remdesivir are being investigated clinically.

SARS-CoV-2 utilizes an envelope homotrimeric Spike glycoprotein (S) to interact with cellular receptor ACE2 ^2,13,14^. Binding with ACE2 triggers a cascade of cell membrane fusion events for viral entry. Each S protomer consists of two subunits: a globular S1 domain at the N-terminal region, and the membrane-proximal S2 and transmembrane domains. Determinants of host range and cellular tropism are found in the RBD within the S1 domain, while mediators of membrane fusion have been identified within the S2 domain ^15,16^. We and others have recently determined the high-resolution structure of SARS-CoV-2 RBD bound to the N-terminal peptidase domain of ACE2 ^14,17^. The overall ACE2-binding mechanism is virtually the same between SARS-CoV-2 and SARS-CoV RBDs, indicating convergent ACE2-binding evolution between these two viruses ^18-22^. This suggests that disruption of the RBD and ACE2 interaction would block SARS-CoV-2 entry into the target cell. Indeed, a few such disruptive agents targeted to ACE2 have been shown to inhibit SARS-CoV infection ^23,24^. However, given the important physiological roles of ACE2 *in viv*o ^25^, these agents may have undesired side effects. Anti-RBD antibodies, on the other hand, are therefore more favorable. Furthermore, SARS-CoV or MERS-CoV RBD-based vaccine studies in experimental animals have also shown strong polyclonal antibody responses that inhibit viral entry ^15,26^. Such critical proof-of-concept findings indicate that anti-RBD antibodies should be able to effectively block SARS-CoV-2 entry. Here, we report on the isolation and characterization of 206 RBD-specific mAbs derived from single B cells of eight SARS-CoV-2 infected individuals. Using bioinformatic and biologic characterization, we identified several mAbs with potent neutralizing activity against pseudovirus and live SARS-CoV-2. However, no cross-activity between RBDs from SARS-CoV or MERS-CoV was found, suggesting that the RBD-based antibody response is viral species-specific. The potent neutralizing antibodies identified here are promising candidates for prophylactic and therapeutic SARS-CoV-2 interventions.

## Results

### Plasma and B cell responses specific to SARS-CoV-2

We collected cross-sectional and longitudinal blood samples from eight SARS-CoV-2-infected subjects during the early outbreak in Shenzhen (Table S1). Samples were named by patient number and either A, B, or C depending on collection sequence. Six patients (P#1 through P#4, P#8, and P#16) had Wuhan travel history and the remaining two (P#5 and P#22) had direct contact with those from Wuhan. P#1 through P#5 is a family cluster with the first documented case of human-to-human transmission of SARS-CoV-2 in Shenzhen ^5^. All subjects recovered and were discharged from the hospital except for P#1 who succumbed to disease despite intensive intervention. To analyze antibody binding, serial plasma dilutions were applied to enzyme-linked immunosorbent assay (ELISA) plates coated with either recombinant RBD or trimeric Spike derived from SARS-CoV-2, SARS-CoV, and MERS-CoV or recombinant NP from SARS-CoV-2. Binding activity was visualized using anti-human IgG secondary antibodies at an optical density (OD) of 450nm. Varying degrees of binding were found across individuals and among samples from the same individual. Samples from P#1, P#2, P#5, and P#16 demonstrated higher binding to both SARS-CoV-2 RBD and NP than the rest (Figure 1A). Three sequential plasma samples collected from P#2 over nine days during early infection showed similar binding to SARS-CoV-2 RBD and NP and remained relative stable over the course of the infection. To our surprise, virtually no cross-reactivity between SARS-CoV RBD and MERS-CoV RBD was detected (Figure 1A), despite strong recognition by the positive control antibodies (data not shown). However, strong cross-reactivity was detected against trimeric Spikes from SARS-CoV and MERS-CoV in both ELISA (Figure 1B) and cell-surface staining (Figure S1). All samples except P#4A demonstrated significant levels of cross-binding to SARS-CoV trimeric Spike while only those from P#1, P#2 and P#4B cross recognized MERS-CoV trimeric Spike (Figure 1B). None of the plasma samples were reactive to HIV-1 envelope trimer derived from strain BG505 ^27^. The same plasma samples were also evaluated for neutralization of pseudoviruses bearing the Spike proteins of either SARS-CoV-2, SARS-CoV, or MERS-CoV. Consistent with the antibody binding results, varying degrees of neutralizing activities against SARS-CoV-2 were found across individuals (Figure 1C). However, cross-neutralizing against SARS-CoV and MERS-CoV is rather minimal as all plasma samples tested, including healthy control plasma, had negligible levels of neutralization (Figure 1C). No detectable neutralization was found for any plasma sample against the pseudovirus control bearing the HIV-1 envelope MG04 (Figure 1C). Taken together, these results suggest that RBDs from SARS-CoV-2, SARS-CoV, and MERS-CoV are likely to be immunologically distinct despite substantial sequence and structural similarities ^14,17^. Thus, regions beyond RBDs likely contribute to the observed cross-reactivity against SARS-CoV and MERS-CoV Spike protein.

**Figure 1.**
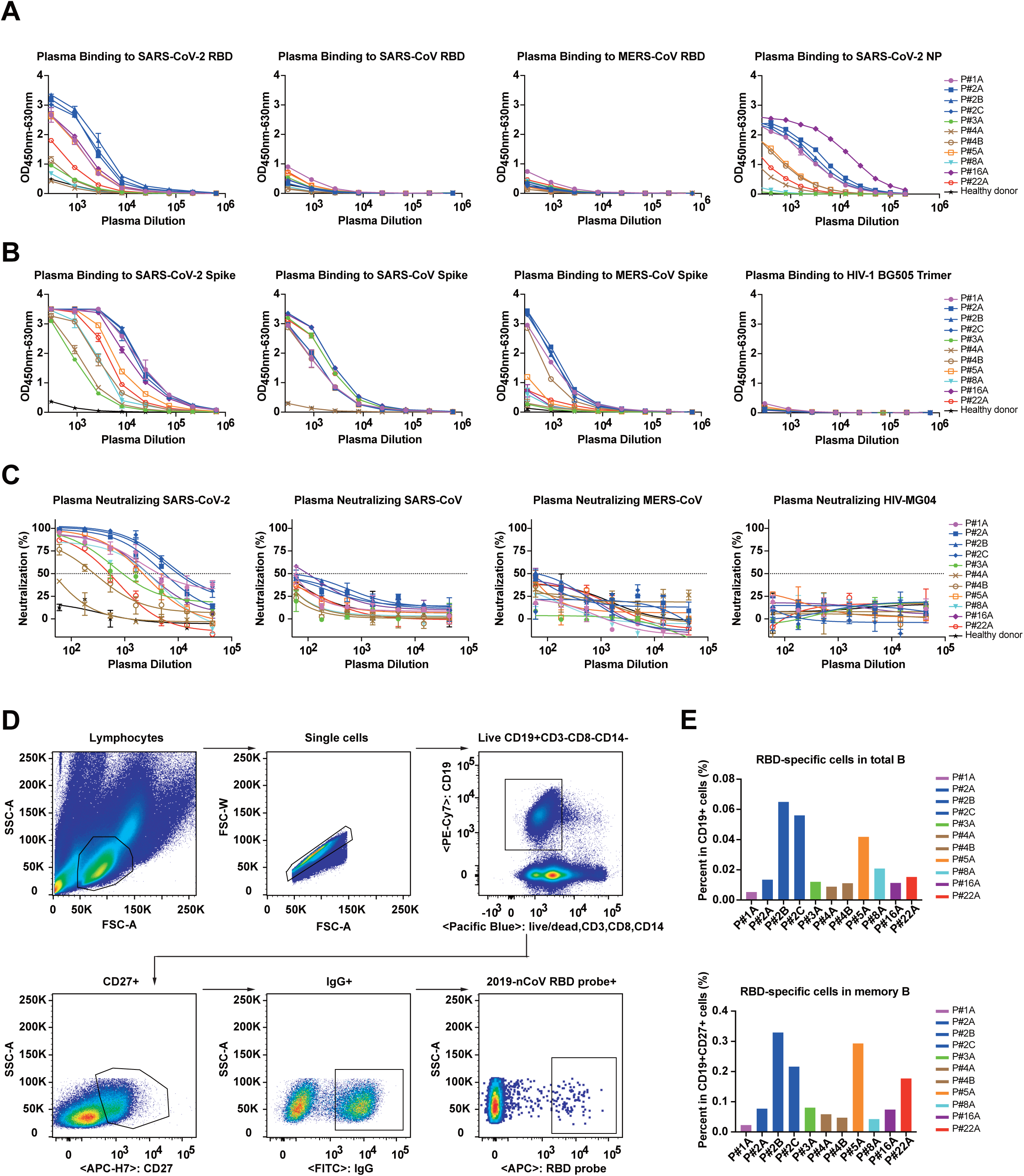
Analyses of plasma and B cell responses specific to SARS-CoV-2. Serial dilutions of plasma samples were analyzed for binding to the (A) RBDs or (B) trimeric Spikes of SARS-CoV-2, SARS-CoV and MERS-CoV by ELISA and (C) for neutralizing activity against pseudoviruses bearing envelope glycoprotein of SARS-CoV-2, SARS-CoV and MERS-CoV. Binding to SARS-CoV-2 NP protein was also evaluated (A). All results were derived from at least two independent experiments. (D) Gating strategy for analysis and isolation of RBD-specific memory B cells and (E) their representation among the total and memory subpopulation of B cells in the eight study subjects. Samples were named as either A, B, or C depending on collection sequence. FSC-W, forward scatter width; FSC-A, forward scatter area; and SSC-A side scatter area.

Flow cytometry with a range of gating strategies was used to study SARS-CoV-2-specific B cell responses and identity B cells recognizing fluorescent-labeled RBD probes (Figure 1D and Figure S2). As shown in Figure 1E, the RBD-specific B cells constitute about 0.005-0.065% among the total B cell population and 0.023-0.329% among the memory subpopulations. The number of RBD-specific B cells are relatively higher in P#2, P#5, P#16, and P#22 (Figure 1E), which appeared to correlate well with binding activity of corresponding plasma samples to SARS-CoV-2 RBD and trimeric Spike protein (Figure 1A and 1B). However, sample P#1A demonstrated the lowest RBD-specific B cell response despite high-level plasma binding. As P#1 was the only patient succumb to disease, it is uncertain whether this dichotomy of high plasma binding activity and low levels of RBD-specific B cells is a surrogate marker of rapid disease progression. This phenomenon needs study in a larger population of samples.

### Single B cell antibody cloning and heavy chain repertoire analyses

We further isolated RBD-binding B cells into single cell suspension for cloning and evaluation of the mAb response (Figure 1D and Figure S2). Immunoglobulin heavy and light chains were amplified by RT-PCR using nested primers. The amplified products were cloned into linear expression cassettes to produce full IgG1 antibodies as previously described ^28,29^. The number of B cell clones varied from 10 to 106 among the subjects and each clone has been differentially represented (Figure S3). Individual IgGs were produced by transfection of linear expression cassettes and tested for SARS-CoV-2 RBD reactivity by ELISA. On average, fifty-eight percent of the antibody clones were reactive, although great variability was found among different individuals (Figure S3). Out of 358 antibodies, we obtained 206 that bound to SARS-CoV-2 RBD with 165 distinct sequences (Table S2). These 206 antibodies demonstrated significant differences in binding activity. For example, a large number of antibodies from samples P#2B, P#2C, P#4A, P4#B, P#5A, P#16A, and P#22A had OD 450 values well over 4.0, while none of those from sample P#1A exceeded 4.0. There were too few antibodies from P#3A and P#8A to make meaningful evaluations (Figure S3). Furthermore, samples from different study subjects also demonstrated substantial differences in heavy chain variable gene (VH) usage (Figure 2A). For instance, P#1 samples are dominated by VH3-53, 3-13, and 1-69 which constituted approximately 21.4%, 14.3%, and 14.3% of the entire VH repertoire, respectively. Samples from P#2 and P#5 are more diverse in distribution and frequency of their VH usage. However, no single or group of VH families stood out among study subjects, suggesting patients have immunologically distinct responses to SARS-CoV-2 infection. This hypothesis is supported by the phylogenetic analysis of all 206 VH sequences superimposed with their corresponding binding activities as presented in Figure 2B. The high-binding clusters (light orange circle: 80% of clusters with OD 450 > 3) were widely distributed across multiple heavy chain families. In fact, majority of the high-binding antibodies were derived by clonal expansion of specific VH families in P#2, P#4, and P#5. Similarly, the middle-(60-80% of clusters with OD 450 > 3) and low- (< 60% cluster with OD 450 > 3) binding clusters were also widely distributed and each consisted of disproportionally represented VH gene families.

**Figure 2.**
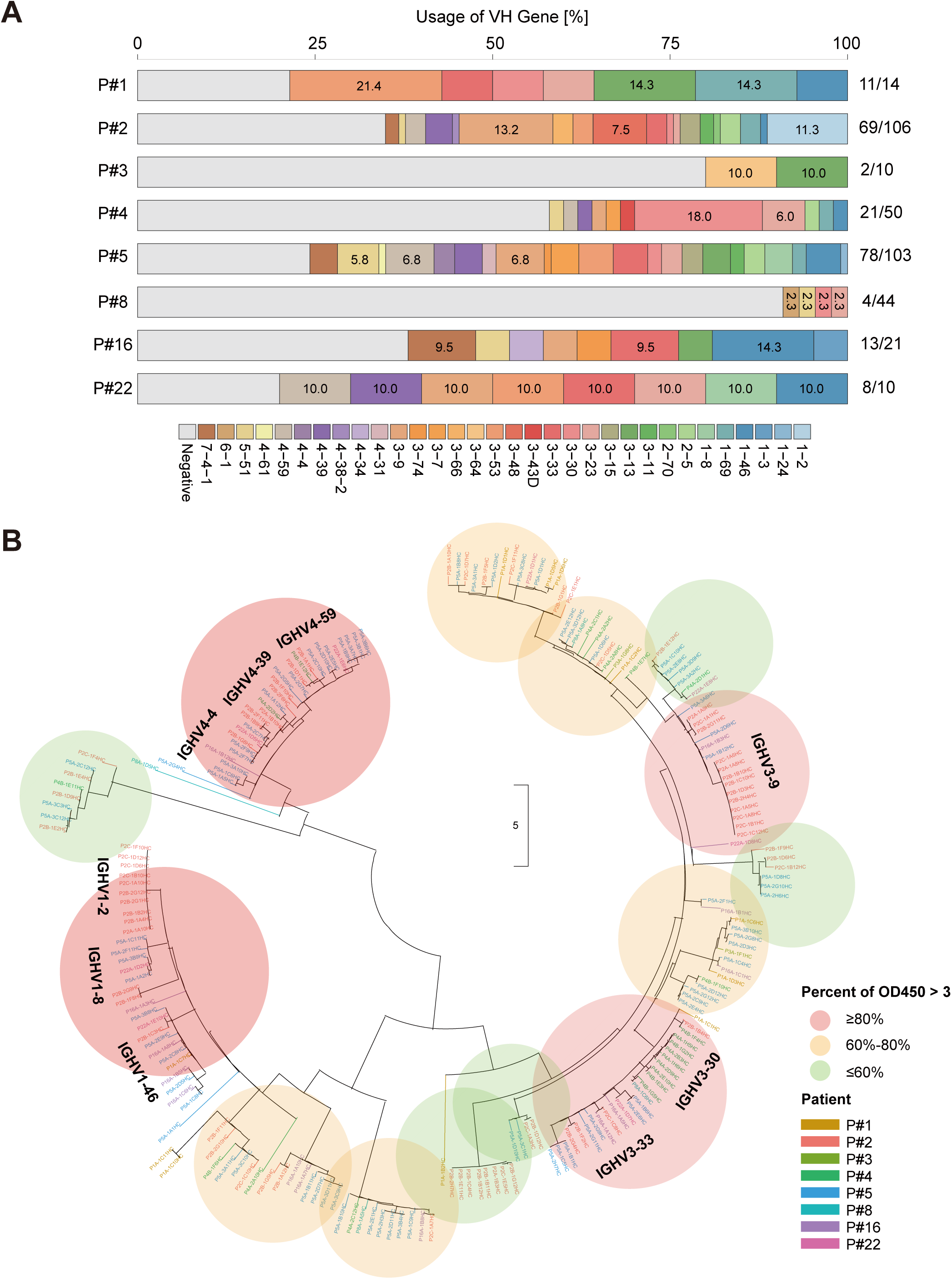
Heavy chain repertoires of SARS-CoV-2 RBD-specific antibodies analyzed (A) by individual subject or (B) across the eight subjects. (A) Distribution and frequency of heavy chain variable (VH) genes usage in each subject shown along the horizontal bar. The same color scheme is used for each VH family across all study subjects. The VHs that dominate across isolated antibodies are indicated by actual frequencies in their respective color boxes. The number of RBD-binding antibodies versus total antibodies isolated are shown on the right. (B) Clustering of VH genes and their association with ELISA binding activity across the eight subjects. Unrooted phylogenetic tree depicting the genetic relationships among all VH genes of the RBD-binding antibodies. Branch lengths are drawn to scale so that sequence relatedness can be readily assessed. Sequences from the same study subject are shown in the same color at the branch tips. Colored circles represent the proportion (light orange, > 80%; light yellow, 60%-80%; light green < 60%) of VH clusters that bind to SARS-CoV-2 RBD with OD 450 values larger than 3. The VH gene families for the highest binding clusters are shown.

### Broad diversity and clonal expansion of antibodies in the repertoire

As P#2 showed a large number of RBD-binding antibodies and was the only patient with three sequential blood samples, we conducted more detailed characterization of P#2 antibodies. Among a total of 69 antibodies from P#2, the majority (59%) were scattered across various branches and the remaining (41%) were clonally expanded into three major clusters (Figure 3A). Antibodies from the three time points (A, B, C) do not appear to group together but rather interdigitate among themselves, suggesting they are highly related during early infection. Three clones were significantly enriched and each constituted between 12-14% of the entire tested repertoire (Figure 3A). Their heavy-chain variable regions belong to the VH1-2*06, VH3-48*02, and VH3-9*01 families. The corresponding light-chain kappa (Igk) belongs to 2-40*01/2D-40*01, 3-20*01, and light-chain lambda (Igl) to 2-14*02 with the respective joining segment kappa 4 (Jk4), Jk5 and joining segment lambda 1 (Jl1) (Table S2). More importantly, these clonally expanded antibodies were identified in all three samples indicating that they are strongly selected for during infection. When comparing their representation within each cluster, VH1-2*06 and VH3-9*01 appeared to increase from approximately 33 to 45%, whereas VH3-48*02 decreased from 33 to 9% over the three time points, although the number of clones was too small for statistical significance. Interestingly, the somatic hypermutation (SHM) or germline divergence for VH1-2*06 was 0% and this cluster persisted during the study period. However, the SHM for VH3-48*02 reached as high as 9.6% and for VH3-9*01 reached 3.8% compared to the overall average of 2.2% ± 3.3 % among the 69 VH sequences. Furthermore, the CDR3 length for VH1-2*06, VH3-48*02, and VH3-9*01 was 19aa, 16aa, and 23aa, respectively, compared with the overall average of 16 ± 4aa among the 69 VH sequences. Close examination of the longest CDR3 from the VH3-9*01 cluster revealed richness in tyrosine, indicating potential hydrogen bonding and hydrophobic interactions with the surrounding residues. These results shed light on the clonal expansion and broad diversity of RBD-specific antibodies during early infection and their potential role in controlling SARS-CoV-2 infection.

**Figure 3.**
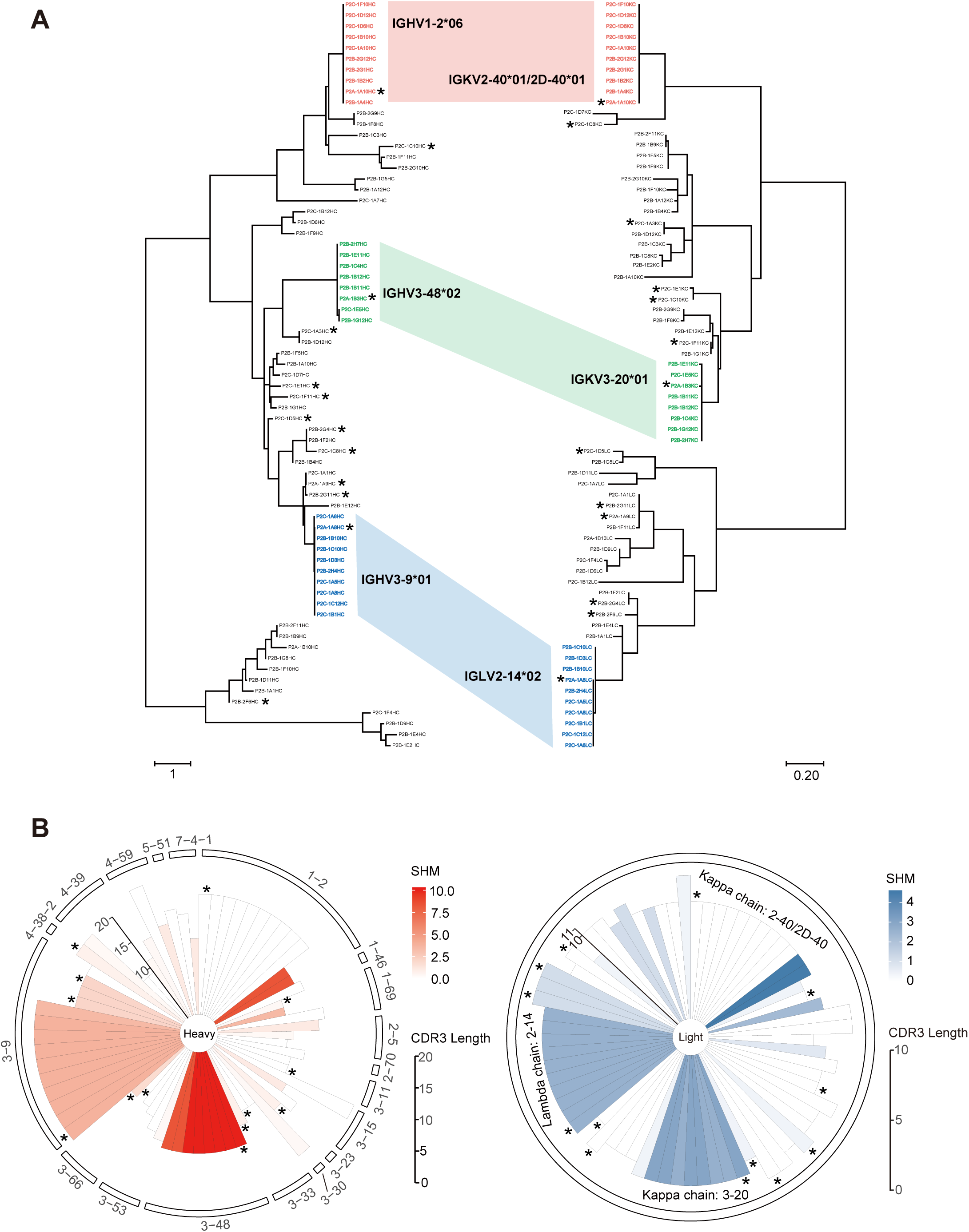
Clonal expansion of specific heavy and light chain families in the P#2 antibody repertoire. (A) Phylogenetic analysis of VH (left) and VL (right) genes for all RBD-binding antibodies. Clonal expanded VH and VL clusters are paired and highlighted in three different colors. Branch lengths are drawn to scale so that sequence relatedness can be readily assessed. (B) Clonal expansion in relation to members of other VH and VL families based on somatic hypermutations (SHM) and CDR3 loop lengths. For the pie charts of VH (left) and VL (right) genes, the radii represent the CDR3 loop length and the color scale indicates the degree of SHM. Heavy and light chain repertoires for each antibody are shown along the pie circles.

### Binding and neutralizing properties of selected antibodies

We selected 13 of the 69 P#2 antibodies sequences based on their representation and distribution on the phylogenetic tree (Figure 3A, starred). Five P#1A antibody clones were used as controls. Surface plasmon resonance (SPR) with SARS-CoV-2 RBD showed that P#2 antibodies had dissociation constants (Kd) ranging from 10^−8^ to 10^−9^ M while those from P#1 ranged from not detectable to 10^−9^ M (Table 1 and Figure S4). SHM did not appear to correlate with Kd; some germline clones with 0% divergence in both VH and VL genes (P2A-1A10, P2B-2G4, P2C-1A3, and P2C-1E1) had Kd values comparable to clones with higher levels of SHM. The Kd of representative clones (P2A-1A8, P2A-1A10, and P2A-1B3) from the three clonally expanded clusters fell into a similar range, suggesting that their expansion may not be driven by affinity maturation. Next, we measured each antibody for competition with ACE2 for binding to the SARS-CoV-2 RBD. Specifically, the RBD was covalently immobilized on a CM5 sensor chip and first saturated by antibody and then flowed through with soluble ACE2. Competing capacity of each antibody was measured as percent reduction in ACE2 binding with the RBD (Table 1 and Figure S5). As shown in Table 1, the evaluated antibodies demonstrated various competing capacity with ACE2. The most powerful were P2C-1F11 and P2B-2F6, which reduced ACE2 binding about 99.2% and 98.5%, respectively. Two of the three representative antibodies from the clonal expanded clusters (P2A-1A10 and P2A-1B3) had slightly over 80% and 90% reduction, respectively. The third representative (P2A-1A8) only showed 57% reduction. Many antibodies had only limited competing power with ACE2 despite impressive Kd values, suggesting binding affinity is not predictive of ACE2 competing capacity. Control antibodies from P#1 demonstrated even lower competing power with ACE2. Surprisingly, none of the antibodies tested demonstrated cross-binding with SARS-CoV and MERS-CoV RBD except P1A-1C7 (Kd=4.85μM), for which only limited cross reactivity with SARS-CoV RBD was detected (Figure S4).

**Table 1.** Binding capacity, neutralizing activity, and heavy chain gene family analysis of 18 monoclonal Abs isolated from Patient #1 and Patient #2.

| Patient | mAbs | Binding to RBD |  | Pseudovirus neutralization |  | Gene family analysis |  |  |  |  |
| --- | --- | --- | --- | --- | --- | --- | --- | --- | --- | --- |
|  |  | Kd (nM) | competing w/ ACE2 | IC<sub>50</sub> (μg/ml) | IC<sub>80</sub> (μg/ml) | IGHV | IGHJ | IGHD | CDR3 length | SHM (%) |
| P#1 | P1A-1C7 | 51.08 | -5.88 | >50 | >50 | 1-46*01,1-46*03 | 4*02 | 2-2*01 | 15 | 0.00 |
|  | P1A-1C10 | 8.48 | 24.51 | >50 | >50 | 1-69*09 | 4*02 | 3-3*01 | 16 | 10.42 |
|  | P1A-1B2 | n.d. | n.d. | n.d. | n.d. | 3-30*03,3-30*18,3-30-5*01 | 4*02 | 5-24*01 | 12 | 11.46 |
|  | P1A-1C1 | n.d. | n.d. | n.d. | n.d. | 3-33*01,3-33*05,3-33*06 | 4*02 | 3-10*01 | 17 | 6.25 |
|  | P1A-1D1 | 260.50 | 6.20 | >50 | >50 | 3-53*01 | 4*02 | 6-13*01 | 12 | 4.21 |
| P#2 | P2A-1A10 | 4.65 | 80.65 | 8.57 | 39.44 | 1-2*06 | 2*01 | 2-2*01 | 19 | 0.00 |
|  | P2C-1C10 | 15.23 | 71.17 | 2.62 | 4.64 | 1-69*01,1-69D*01 | 4*02 | 4-23*01 | 11 | 0.35 |
|  | P2C-1A3 | 2.47 | 81.21 | 0.62 | 5.94 | 3-11*04 | 5*01,5*02 | 6-13*01 | 12 | 0.00 |
|  | P2C-1D5 | 1.64 | 17.61 | 10.65 | 25.36 | 3-23*04 | 4*02 | 3-10*01 | 14 | 0.69 |
|  | P2B-2G4 | 21.29 | 41.03 | 5.11 | >50 | 3-33*01,3-33*06 | 4*02 | 5-18*01 | 11 | 0.00 |
|  | P2C-1C8 | 8.76 | 74.45 | 34.38 | >50 | 3-33*01,3-33*06 | 4*02 | 3-22*01 | 13 | 0.69 |
|  | P2A-1B3 | 6.00 | 92.15 | 16.77 | >50 | 3-48*02 | 5*02 | 3-10*01 | 16 | 10.07 |
|  | P2C-1E1 | 14.99 | 56.61 | >50 | >50 | 3-66*01,3-66*04 | 4*02 | 5-12*01 | 9 | 0.00 |
|  | P2C-1F11 | 2.12 | 99.17 | 0.03 | 0.12 | 3-66*01,3-66*04 | 6*02 | 2-15*01 | 11 | 1.75 |
|  | P2A-1A8 | 8.91 | 57.06 | 7.68 | 26.41 | 3-9*01 | 6*02 | 5-12*01 | 23 | 3.82 |
|  | P2A-1A9 | 15.18 | 53.60 | 26.27 | >50 | 3-9*01 | 6*02 | 3-22*01 | 17 | 2.08 |
|  | P2B-2G11 | 17.57 | 52.28 | 34.84 | >50 | 3-9*01 | 6*02 | 1-26*01 | 17 | 2.08 |
|  | P2B-2F6 | 5.14 | 98.50 | 0.05 | 0.61 | 4-38-2*02 | 3*02 | 2-2*01 | 20 | 0.69 |

The program IMGTV-QUEST was applied to analyze gene germline, complementarity determining region (CDR) 3 length, and somatic hypermutation (SHM). The CDR3 length was calculated from amino acids sequences. The SHM frequency was calculated from the mutated nucleotides. n.d.: not detectable.

We next studied antibody neutralizing activities against pseudoviruses bearing the Spike protein of SARS-CoV-2. Consistent with the competing capacity findings, neutralizing activity varied considerably with IC_50_ values ranging from 0.03 to > 50 μg/ml (Figure 4B, 4A and Table 1). P2C-1F11 and P2B-2F6 were the most potent, followed by P2C-1A3 and P2C-1C10. Overall, ACE2 competing capacity correlated well with the neutralizing activities, although this correlation was not exact in some instances. Notably, no cross-neutralization was found either against pseudoviruses bearing the Spike of SARS-CoV or MERS-CoV (data not shown) nor with cell-surface staining of trimeric SARS-CoV and MERS-CoV Spike (Figure S6). Furthermore, we selected P2C-1F11, P2B-2F6, and P2C-1A3 for neutralizing activity analyses against live SARS-CoV-2. Consistent with their respective pseudovirus assay findings, P2C-1F11 and P2B-2F6 demonstrated potent neutralization activity while that of P2C-1A3 was somewhat lower, although it needs to be noted that CPE assay is not particularly quantitative (Figure 4C). Lastly, we determined whether these antibodies compete for similar epitopes on the SARS-CoV-2 RBD. We selected a total of six antibodies with ACE2 competitive capacities of at least 70% and analyzed them in a pairwise competition fashion using SPR. As shown in Table 2 and Figure S7, variable degrees of competition were found among the pairs of antibodies. P2C-1A3, for instance, was competitive against all antibodies tested with reduction capacity ranging from 52 to 76. P2C-1F11, on the other hand, was less competitive with other antibodies and in particular, only minimally competitive with P2C-1C10. P2B-2F6, another potent neutralizing antibody, was broadly competitive with all antibodies tested. These results indicate that the antibodies analyzed recognized both overlapping and distinct epitopes. Different mAbs may therefore exert their neutralizing activity through different mechanisms.

**Table 2.** Epitope mapping of mAbs through competitive binding to SARS-CoV-2 RBD.

| mAbs | P2C-1A3 | P2C-1C10 | P2C-1F11 | P2B-2F6 | P2A-1B3 | P2A-1A10 |
| --- | --- | --- | --- | --- | --- | --- |
| P2C-1A3 |  | 68.41 | 57.44 | 76.73 | 65.20 | n.a. |
| P2C-1C10 | 75.33 |  | -0.32 | 49.69 | 42.98 | n.a. |
| P2C-1F11 | 52.05 | -2.31 |  | 42.65 | 5.98 | n.a. |
| P2B-2F6 | 74.87 | 70.97 | 30.22 |  | 52.79 | n.a. |
| P2A-1B3 | 57.94 | 63.83 | 14.35 | 51.88 |  | n.a. |
| P2A-1A10 | 76.31 | 84.27 | 79.50 | 73.92 | 42.19 |  |

n.a.: not applicable

**Figure 4.**
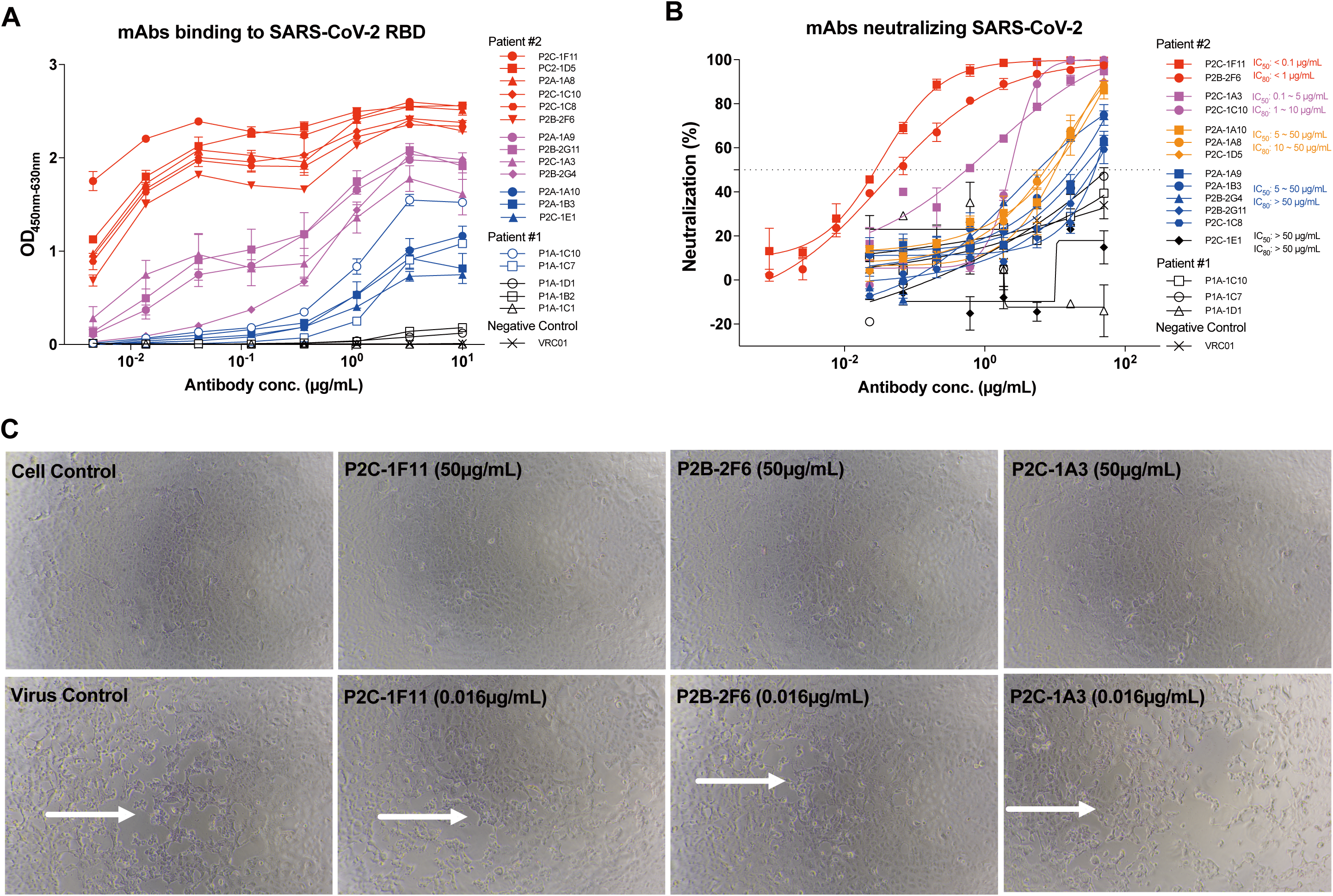
Antibody neutralization analyzed by pseudovirus and live SARS-CoV-2. (A) Quality control of antibody through ELISA analysis prior to neutralization assay. A serial dilution of each antibody was evaluated against SARS-CoV-2 RBD coated on the ELISA plate and their binding activity was recorded at an optical density (OD) of 450nm and 630nm. (B-C) Antibody neutralization analyzed by pseudovirus (B) or live SARS-CoV-2 (C). A serial dilution of each antibody was tested against pseudovirus while two dilutions against live SARS-CoV-2. Cytopathic effects (CPE) were observed daily and recorded on Day 2 post-exposure. Selected antibodies and their concentrations tested are indicated at the upper left corner.

## Discussion

We characterized antibody responses in eight COVID-19 patients and isolated 206 mAbs specific to the SARS-CoV-2 RBD. Bioinformatic and biologic characterization indicates that these antibodies are derived from broad and diverse families of antibody heavy and light chains. Each individual appears to have unique pattern of distribution in the antibody repertoire without apparent preferences for particular antibody families. Each antibody clone is also differentially represented. In P#2, for whom additional analyses were conducted, we found substantial variability in the distribution and frequency of each antibody family. Some clones were identified only once whereas others underwent high degrees of clonal expansion. Some clones were virtually identical to their germline ancestors while others became more divergent during the infection period. The CDR3 length also varied among the different clones. These differences at the genetic levels corresponded with their binding and neutralizing activities. Binding affinity (Kd) fell in the range of 10^−8^ to 10^−9^ M, equivalent to many antibodies identified during acute infections ^30-32^ but significantly lower than those identified during chronic HIV-1 infections ^33-35^. However, binding affinity alone does not predict neutralizing activity. Competition with the receptor ACE2 governs antibody potency, although some degree of discrepancy does exist. In particular, the most potent antibodies, P2C-1F11 and P2B-2F6, out-competed ACE2 with close to 100% efficiency, indicating that blocking the RBD and ACE2 interaction is a useful surrogate for antibody neutralization. Among the antibodies tested, substantial variations in competition for similar RBD epitopes or regions were also found. The most potent antibody, P2C-1F11, did not seem target the same epitope as the relatively moderate antibody P2C-1C10. Thus, these two antibodies could be combined for synergistic antiviral effect. As we continue to screen more antibodies from P#2 and other study subjects, more potent and diverse antibodies are expected to be identified. These antibodies will serve as the best candidates for the development of prophylactic and therapeutic intervention against COVID-19 infection.

Most surprising in this study was the absence of antibody cross-reactivity with RBDs from SARS-CoV and MERS-CoV. Based on the sequential and structural similarities of RBDs from SARS-CoV-2 and SARS-CoV, we predicted some degree of cross-binding and even cross-neutralization between the two viruses. However, species-specific RBD responses in SARS-CoV-2 patients do suggest that RBDs from SARS-CoV-2 and SARS-CoV are immunologically distinct. If so, antibodies and vaccines must target each viral species differently in order to achieve maximum efficacy in protecting the host from infection. Our finding somewhat resolves the question of why many previously isolated SARS-CoV antibodies failed to cross-neutralize SARS-CoV-2 despite detectable levels of binding with Spike of SARS-CoV-2^36^. The absence of cross-recognition between RBDs was also apparent at the plasma level. Although strong binding to SARS-CoV-2 RBD was identified, plasma samples from the study subjects failed to demonstrate appreciable cross-reactivity with either SARS-CoV or MERS-CoV RBD, highlighting the immunological distinctions among the RBDs from the three viruses. However, substantial cross-reactivity were found when the same plasma samples were applied to the trimeric Spike proteins of SARS-CoV and MERS-CoV, although this was higher with the former than the latter. This indicates that such cross-reactivity likely occurs in regions outside the RBD. Determining whether this cross-reactive response has any neutralizing or protection capacity against infection would require further investigation. Finally, despite successfully isolating and characterizing a large of number mAbs against SARS-CoV-2, we cannot draw any firm correlation between antibody response and disease status at this time. In particular, the three severe cases (P#1, P#2, and P#5) appear to have relatively higher plasma binding and neutralizing activities against SARS-CoV-2 than those with relative mild symptoms. A larger number of patients must be studied to elucidate the drivers and impact of associations between antibody response and disease progression, which will provide pivotal reference for our antibody-based intervention as well as vaccine development.

## Materials and Methods

### Study approval

This study received approval from the Research Ethics Committee of Shenzhen Third People’s Hospital, China (approval number: 2020-084). The Research Ethics Committee waived the requirement informed consent before the study started because of the urgent need to collect epidemiological and clinical data. We analyzed all the data anonymously.

### Patients and blood samples

The study enrolled a total of eight patients aged 10 to 66 years old infected with SARS-CoV-2 in January 2020 (Table S1). A plasma sample from a healthy control was also included. Of these eight patients, six (P#1 through P#4, P#8, and P#16) had Wuhan exposure history through personal visit and two had direct contact with individuals from Wuhan. Four subjects (P#1 through P#4) were part of a family cluster (P#1 through P#5) infected while visiting Wuhan and subsequently transmitted infection to P#5 after returning to Shenzhen ^5^. All patients were hospitalized at Shenzhen Third People’s Hospital, the designated city hospital for treatment of COVID-19 infected patients, three to nine days after symptom onset. All patients presented with fever, fatigue, and dry cough and three (P#1, P#2 and P#5) developed severe pneumonia. Four patients (P#1, P#2, P#5, and P#22) were 60 years or older, of which three (P#1, P#2, and P#22) had underlying disease such as hypertension. SARS-CoV-2 infection status was verified by RT-PCR of nasopharyngeal swab and throat swab specimens. No patient had detectable influenza A, B, respiratory syncytial virus (RSV), or adenovirus co-infections. Chest computed tomographic scans showed varying degrees of bilateral lung patchy shadows or opacity. All patients received antiviral and corticosteroid treatments, recovered and were discharged except for P#1, who succumbed to disease in hospital. Single (P#1, P#3, P#5, P#8, P#16, and P#22) or sequential (P#2 and P#4) blood samples were collected during hospitalization and follow-up visits and separated into plasma and peripheral blood mononuclear cells (PBMCs) by Ficoll-Hypaque gradient (GE Healthcare) centrifugation. All plasma samples were heat-inactivated at 56 °C for 1h before being stored at -80 °C. PBMCs were maintained in freezing media and stored in liquid nitrogen until use.

### Recombinant RBDs and trimeric Spike from SARS-CoV-2, SARS-CoV, and MERS-CoV and receptor ACE2

Recombinant RBDs and trimeric Spike for MERS-CoV, SARS-CoV, and SARS-CoV-2 and the N-terminal peptidase domain of human ACE2 (residues Ser19-Asp615) were expressed using the Bac-to-Bac baculovirus system (Invitrogen) as previously described ^18,19,37-39^. SARS-CoV-2 RBD (residues Arg319-Phe541) containing the gp67 secretion signal peptide and a C-terminal 6×His tag was inserted into pFastBac-Dual vectors (Invitrogen) and transformed into DH10Bac component cells. The bacmid was extracted and further transfected into Sf9 cells using Cellfectin II Reagents (Invitrogen). The recombinant viruses were harvested from the transfected supernatant and amplified to generate high-titer virus stock. Viruses were then used to infect Hi5 cells for RBD and trimeric Spike expression. Secreted RBD and trimeric Spike were harvested from the supernatant and purified by gel filtration chromatography as previously reported ^18,19,37-39^.

### ELISA analysis of plasma and antibody binding to RBD, trimeric Spike, and NP proteins

The recombinant RBDs and trimeric Spike derived from SARS-CoV-2, SARS-CoV and MERS-CoV and the SARS-CoV-2 NP protein (Sino Biological, Beijing) were diluted to final concentrations of 0.5 μg/ml or 2μg/ml, then coated onto 96-well plates and incubated at 4°C overnight. Samples were washed with PBS-T (PBS containing 0.05% Tween 20) and blocked with blocking buffer (PBS containing 5% skim milk and 2% BSA) at RT for 1h. Either serially diluted plasma samples or isolated mAbs were added the plates and incubated at 37°C for 1h. Wells were then incubated with secondary anti-human IgG labeled with HRP (ZSGB-BIO, Beijing) and TMB substrate (Kinghawk, Beijing) and optical density (OD) was measured by a spectrophotometer at 450nm and 630nm. The serially diluted plasma from healthy individuals or mAbs against SARS-CoV, MERS-CoV or HIV-1 were used as controls.

### Isolation of RBD-specific single B cells by FACS

RBD-specific single B cells were sorted as previously described ^28,40^. In brief, PBMCs from infected individuals were collected and incubated with an antibody and RBD cocktail for identification of RBD-specific B cells. The cocktail consisted of CD19-PE-Cy7, CD3-Pacific Blue, CD8-Pacific Blue, CD14-Pacific Blue, CD27-APC-H7, IgG-FITC (BD Biosciences) and the recombinant RBD-Strep or RBD-His described above. Three consecutive staining steps were conducted. The first was a LIVE/DEAD Fixable Dead Cell Stain Kit (Invitrogen) in 50µl phosphate-buffered saline (PBS) applied at RT for 20 minutes to exclude dead cells. The second utilized an antibody and RBD cocktail for an additional 30min at 4 °C. The third staining at 4 °C for 30min involved either: Streptavidin-APC (eBioscience) and/or Streptavidin-PE (BD Biosciences) to target the Strep tag of RBD, or anti-his-APC and anti-his-PE antibodies (Abcam) to target the His tag of RBD. The stained cells were washed and resuspended in PBS before being strained through a 70μm cell mesh (BD Biosciences). RBD-specific single B cells were gated as CD19+CD3-CD8-CD14-IgG+RBD+ and sorted into 96-well PCR plates containing 20μl of lysis buffer (5 μl of 5 x first strand buffer, 0.5 μl of RNase out, 1.25 μl of 0.1 M DTT (Invitrogen) per well and 0.0625 μl of Igepal (Sigma). Plates were then snap-frozen on dry ice and stored at -80 °C until RT reaction.

### Single B cell PCR, cloning and expression of mAbs

The IgG heavy and light chain variable genes were amplified by nested PCR and cloned into linear expression cassettes or expression vectors to produce full IgG1 antibodies as previously described ^29,41^. Specifically, all second round PCR primers containing tag sequences were used to produce the linear Ig expression cassettes by overlapping PCR. Separate primer pairs containing the specific restriction enzyme cutting sites (heavy chain, 5’-AgeI/3’-SalI; kappa chain, 5’-AgeI/3’-BsiWI; and lambda chain, 5’-AgeI/3’-XhoI) were used to amplify the cloned PCR products. The PCR products were purified and cloned into the backbone of antibody expression vectors containing the constant regions of human IgG1. Overlapping PCR products of paired heavy and light chain expression cassettes were co-transfected into 293T cells (ATCC) grown in 24-well plates. Antigen-specific ELISA was used to detect the binding capacity of transfected culture supernatants to SARS-CoV-2 RBD. Monoclonal antibodies were produced by transient transfection of 293F cells (Life Technologies) with equal amounts of paired heavy and light chain plasmids. Antibodies in the culture supernatant was purified by affinity chromatography using Protein A beads columns (National Engineering Research Center for Biotechnology, Beijing) according to the manufacturer’s protocol. Concentrations were determined by BCA Protein Assay Kits (Thermo Scientific). SARS-CoV, MERS-CoV, and HIV-1 mAbs were also included as controls. SARS-CoV antibodies (S230 and m396) previously isolated by others ^42^ were synthesized and sequences verified before expression in 293T cells and purification by protein A chromatography. MERS-CoV antibodies (Mab-GD33) were derived from previously reported ^43^. HIV-1 antibody VRC01 was a broadly neutralizing antibody directly isolated from a patient targeting the CD4 binding site of envelope glycoprotein ^40^.

### Antibody binding kinetics, epitope mapping, and competition with receptor ACE2 measured by SPR

The binding kinetics and affinity of mAbs to SARS-CoV-2 RBD were analyzed by SPR (Biacore T200, GE Healthcare). Specifically, purified RBDs were covalently immobilized to a CM5 sensor chip via amine groups in 10mM sodium acetate buffer (pH 5.0) for a final RU around 250. SPR assays were run at a flow rate of 30ml/min in HEPES buffer. The sensograms were fit in a 1:1 binding model with BIA Evaluation software (GE Healthcare). For epitope mapping, two different antibodies were sequentially injected and monitored for binding activity to determine whether the two mAbs recognized separate or closely-situated epitopes. To determine competition with the human ACE2 peptidase domain, SARS-CoV-2 RBD was immobilized to a CM5 sensor chip via amine group for a final RU around 250. Antibodies (1 μM) were injected onto the chip until binding steady-state was reached. ACE2 (2 μM), which was produced and purified as above, was then injected for 60 seconds. Blocking efficacy was determined by comparison of response units with and without prior antibody incubation.

### Analysis of plasma and antibody binding to cell surface expressed trimeric Spike protein

HEK 293T cells were transfected with expression plasmid encoding the full length spike of SARS-CoV-2, SARS-CoV or MERS-CoV and incubated at 37 °C for 36 h. The cells were digested with trypsin and distributed into 96 well plates for the individual staining. Cells were washed twice with 200µl staining buffer (PBS with 2% heated-inactivated FBS) between each following steps. The cells were stained at room temperature for 30 minutes in 100 μl staining buffer with 1:100 dilutions of plasma or 20 μg/ml monoclonal antibodies. The cells were then stained with PE labeled anti-human IgG Fc secondary antibody (Biolegend) at a 1:20 dilution in 50 μl staining buffer at room temperature for 30 minutes. Finally, the cells were re-suspended and analyzed with FACS Calibur instrument (BD Biosciences, USA) and FlowJo 10 software (FlowJo, USA). HEK 293T cells without transfection were also stained as background control. S230 and m396 targeting the RBD of SARS-CoV spike ^42^ and Mab-GD33 targeting the RBD of MERS-CoV spike ^43^ were used as positive primary antibody controls, while VRC01 targeting HIV-1 env ^40^ was used as an irrelevant primary antibody control.

### Neutralization activity of mAbs against pseudovirus and live SARS-CoV-2

SARS-CoV-2, SARS-CoV and MERS-CoV pseudovirus were generated by co-transfection of human immunodeficiency virus backbones expressing firefly luciferase (pNL43R-E-luciferase) and pcDNA3.1 (Invitrogen) expression vectors encoding the respective S proteins into 293T cells (ATCC) ^37,38,44,45^. Viral supernatants were collected 48 h later. Viral titers were measured as luciferase activity in relative light units (Bright-Glo Luciferase Assay Vector System, Promega Biosciences). Control envelope glycoproteins derived from human immunodeficiency virus (HIV)-1 and their corresponding pseudoviruses were produced in the same manner. Control mAbs included VRC01 against HIV-1 ^40^; S230 and m396 against SARS-CoV ^42^; and Merb-GD33 against MERS-CoV ^43^. Neutralization assays were performed by incubating pseudoviruses with serial dilutions of purified mAbs at 37°C for 1h. Huh7 cells (ATCC) (approximately 1.5 × 10^4^ per well) were added in duplicate to the virus-antibody mixture. Half-maximal inhibitory concentrations (IC_50_) of the evaluated mAbs were determined by luciferase activity 48h after exposure to virus-antibody mixture using GraphPad Prism 6 (GraphPad Software Inc.).

All experiments involving live SARS-CoV-2 followed approved Biosafety Level 3 laboratory standard operating procedures. Neutralization assays against live SARS-CoV-2 were conducted using a clinical isolate (Beta/Shenzhen/SZTH-003/2020, EPI_ISL_406594 at GISAID) previously obtained from a nasopharyngeal swab of P#3. The isolate was amplified in Vero cell lines to make working stocks of the virus (1 × 10^5^ PFU/ml). To analyze the mAb neutralizing activities, Vero E6 cells were seeded at 10^4^/well in 96-well culture plates and cultured at 37 °C to form a monolayer. Serial dilutions of mAbs were mixed separately with 100 PFU of SARS-CoV-2, incubated at 37 °C for 1 h, and added to the monolayer of Vero E6 cells in duplicates. Cells either unexposed to the virus or mixed with 100 PFU SARS-CoV-2 were used as negative (uninfected) and positive (infected) controls, respectively. Cytopathic effects (CPE) were observed daily and recorded on Day 2 post-exposure.

### Gene family usage and phylogenetic analysis of mAbs

The program IMGT/V-QUEST (http://www.imgt.org/IMGT_vquest/vquest) was used to analyze germline gene, germline divergence or degree of somatic hypermutation (SHM), the framework region (FR) and the loop length of the complementarity determining region 3 (CDR3) for each antibody clone. The IgG heavy and light chain variable genes were aligned using Clustal W in the BioEdit sequence analysis package (https://bioedit.software.informer.com/7.2/). Phylogenetic analyses were performed by the Maximum Likelihood method using MEGA X (Molecular Evolutionary Genetics Analysis across computing platforms). Several forms of the phylogenetic trees are presented for clarity.

### Antibody production

The production of antibodies was conducted as previously described ^38,46^. The genes encoding the heavy and light chains of isolated antibodies were separately cloned into expression vectors containing IgG1 constant regions and the vectors were transiently transfected into HEK293T or 293F cells using polyethylenimine (PEI) (Sigma). After 72h, the antibodies secreted into the supernatant were collected and captured by protein A Sepharose (GE Healthcare). The bound antibodies were eluted and further purified by gel-filtration chromatography using a Superdex 200 High Performance column (GE Healthcare). The purified antibodies were either used in binding and neutralizing assays.

## Acknowledgments

We acknowledge the work and contribution of all the health providers from Shenzhen Third People’s Hospital. We also thank patients for their active participation. This study was supported by Bill & Melinda Gates Foundation, the Science and Technology Innovation Committee of Shenzhen Municipality (202002073000002), and by Tsinghua University Initiative Scientific Research Program (20201080053). This work is also partially supported by the National Natural Science Foundation Award (81530065), Beijing Municipal Science and Technology Commission (171100000517-001 and -003), Beijing Advanced Innovation Center for Structural Biology at Tsinghua University, the National Key Plan for Scientific Research and Development of China (grant number 2016YFD0500307), Tencent Foundation, Shuidi Foundation, and TH Capital. The funders had no role in study design, data collection, data analysis, data interpretation, or writing of the report.

## Author contributions

LZ, ZZ, LL and SZ conceived and designed the study. BJ and QZ performed most of the experiments together with assistance from XG, RW, JY, SS, BJ, SS, and XS. XT performed live SARS-CoV-2 neutralization assay. JY, JG, JL, XW provided assistance in RBD and trimeric Spike protein production. JY and LL played critical roles in recruitment and clinical management of the study subjects. HW and JZ are in charge of sample collection and processing. YW provides additional pseudovirus assay for measuring neutralizing activity against SARS-CoV-2. BJ, QZ, ZZ and LZ had full access to data in the study, generated figures and tables, and take responsibility for the integrity and accuracy of the data presentation. LZ and ZZ wrote the manuscript. All authors reviewed and approved the final version of the manuscript.

## Data availability statements

We are happy to share reagents and information presented in this study upon request.

## Figure Legends

**Figure S1.**
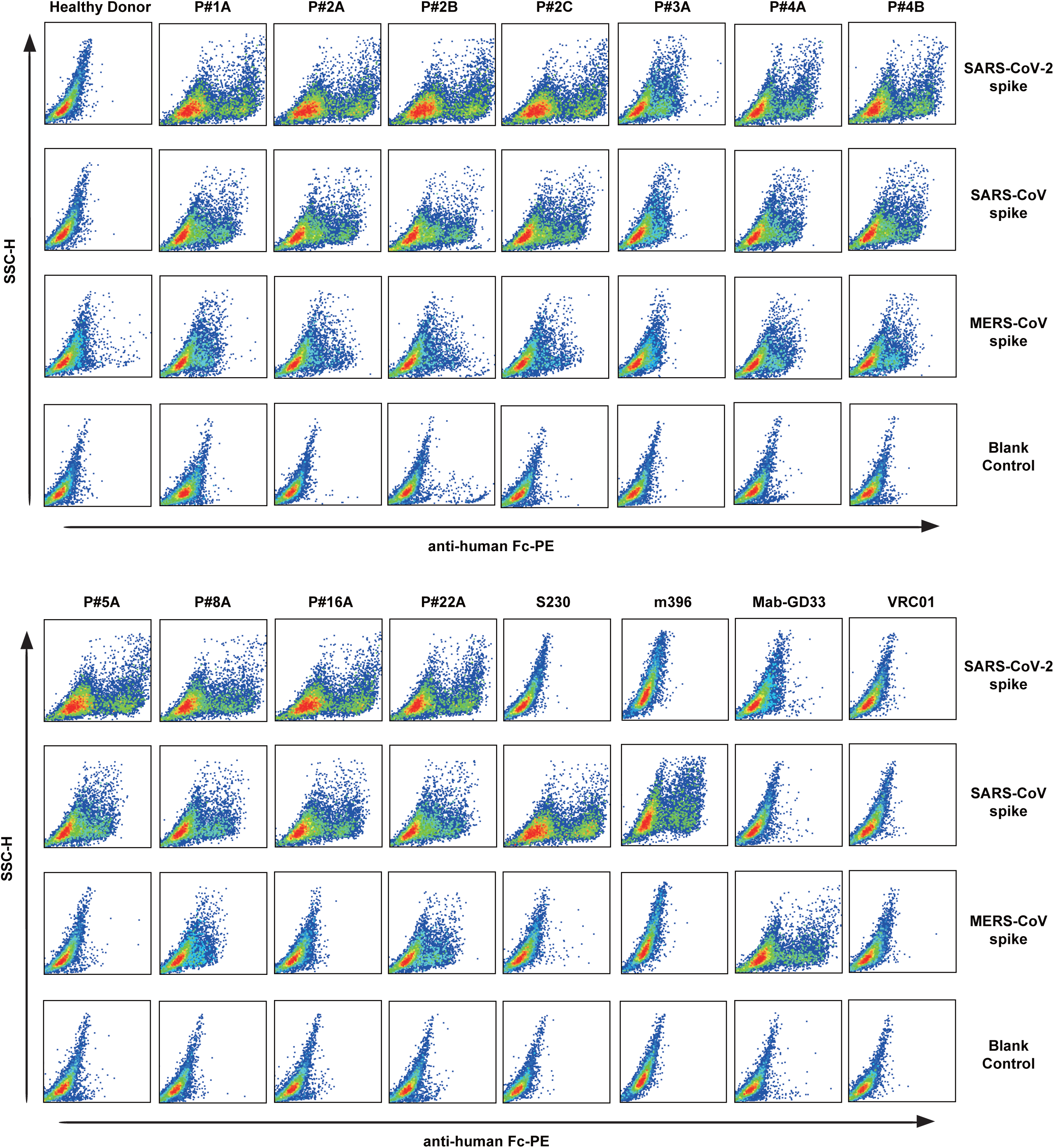
Analysis of plasma binding to cell surface expressed trimeric Spike protein. HEK 293T cells transfected with expression plasmid encoding the full length spike of SARS-CoV-2, SARS-CoV or MERS-CoV were incubated with 1:100 dilutions of plasma from the study subjects. The cells were then stained with PE labeled anti-human IgG Fc secondary antibody and analyzed by FACS. Positive control antibodies include S230 and m396 targeting the RBD of SARS-CoV Spike, and Mab-GD33 targeting the RBD of MERS-CoV Spike. VRC01 is negative control antibody targeting HIV-1 envelope glycoprotein.

**Figure S2.**
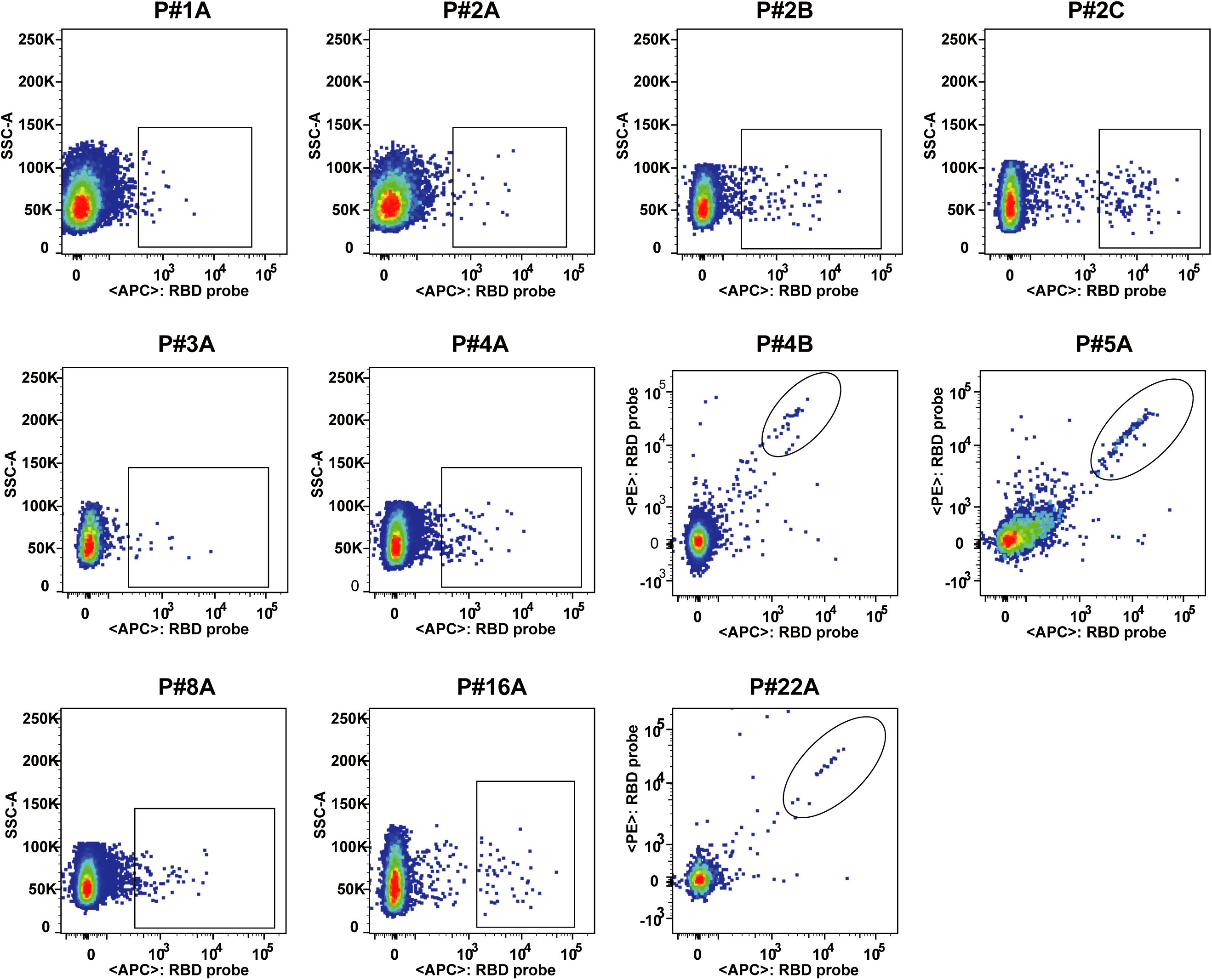
RBD-specific memory B cells analyzed and isolated through FACS. The recombinant RBD was labeled with either a Strep or His tag and used alone or in combination to identify and isolate RBD-specific single B cells through staining with the Streptavidin-APC and/or Streptavidin-PE, or anti-His-APC and anti-His-PE antibodies. B cells to be isolated are highlighted in boxes or ovals. Samples were named as either A, B, or C depending on collection sequence. FSC-W, forward scatter width; FSC-A, forward scatter area; and SSC-A side scatter area.

**Figure S3.**
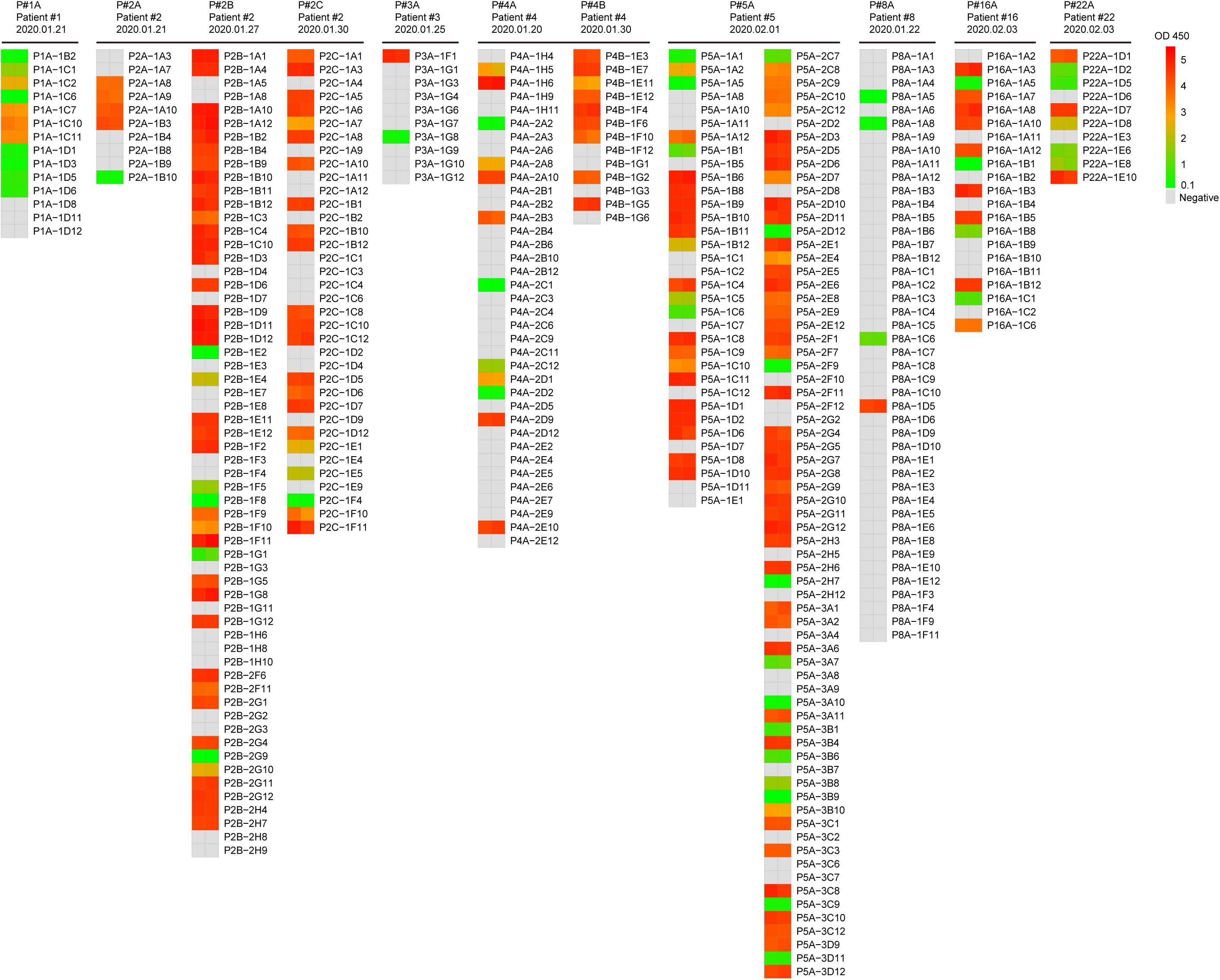
ELISA screening of SARS-CoV-2 RBD-specific antibodies in the supernatant of transfected cells. The study subjects and the date of sampling are indicated on the top. Samples were named as either A, B, or C depending on collection sequence. Antibodies tested for each sample are aligned in one vertical column whenever possible. For each evaluated antibody, at least two independent measurements were performed and are presented adjacently on the same row. Binding activities were assessed by OD 450 and indicated by the color scheme on the right. Negatives (no binding activity) are shown in gray for OD 450 values less than 0.1.

**Figure S4.**
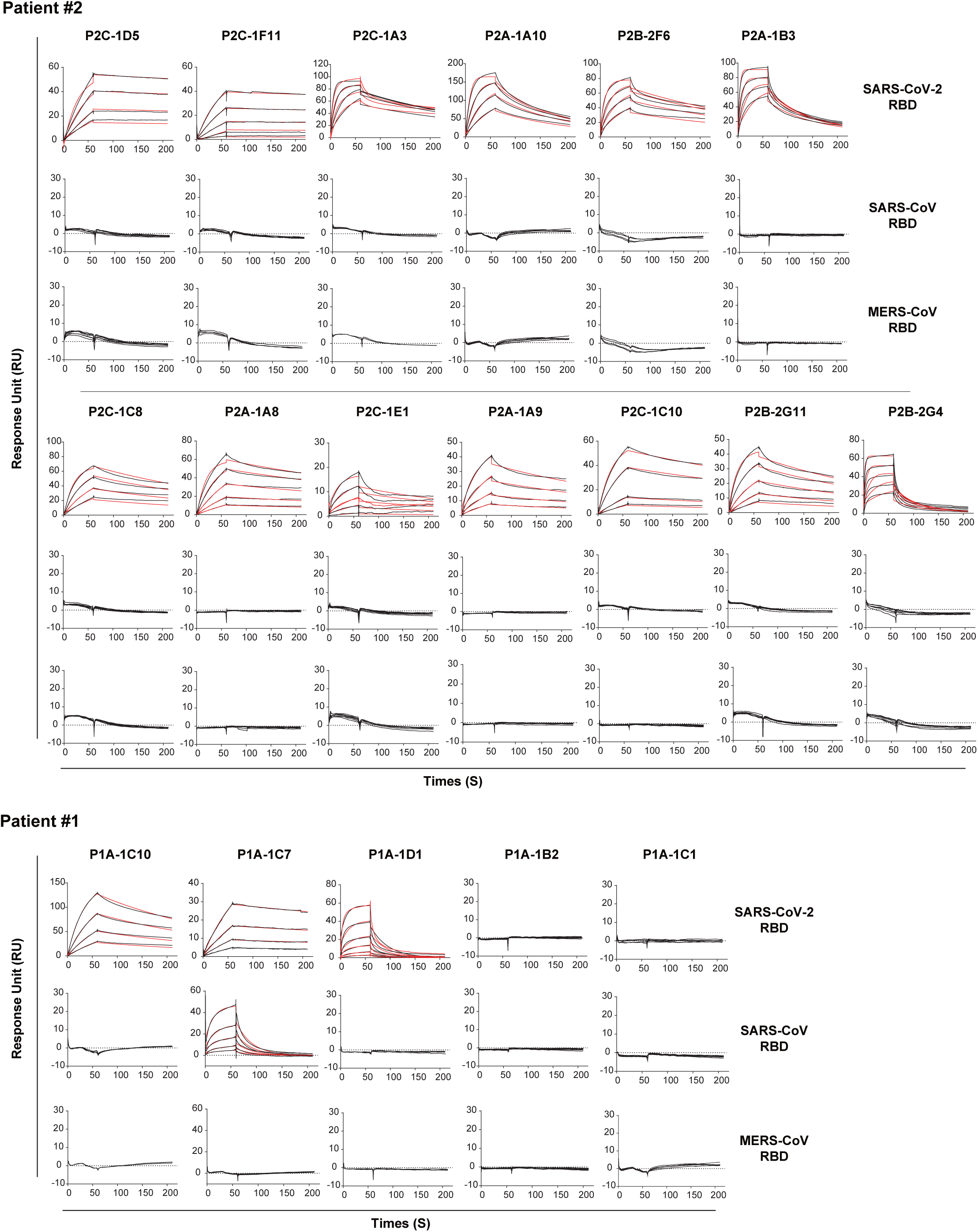
Binding kinetics of isolated mAbs with SARS-CoV-2 RBD measured by SPR. The purified soluble SARS-CoV-2 RBD were covalently immobilized onto a CM5 sensor chip followed by injection of individual antibody at four or five different concentrations. The black lines indicate the experimentally derived curves while the red lines represent fitted curves based on the experimental data.

**Figure S5.**
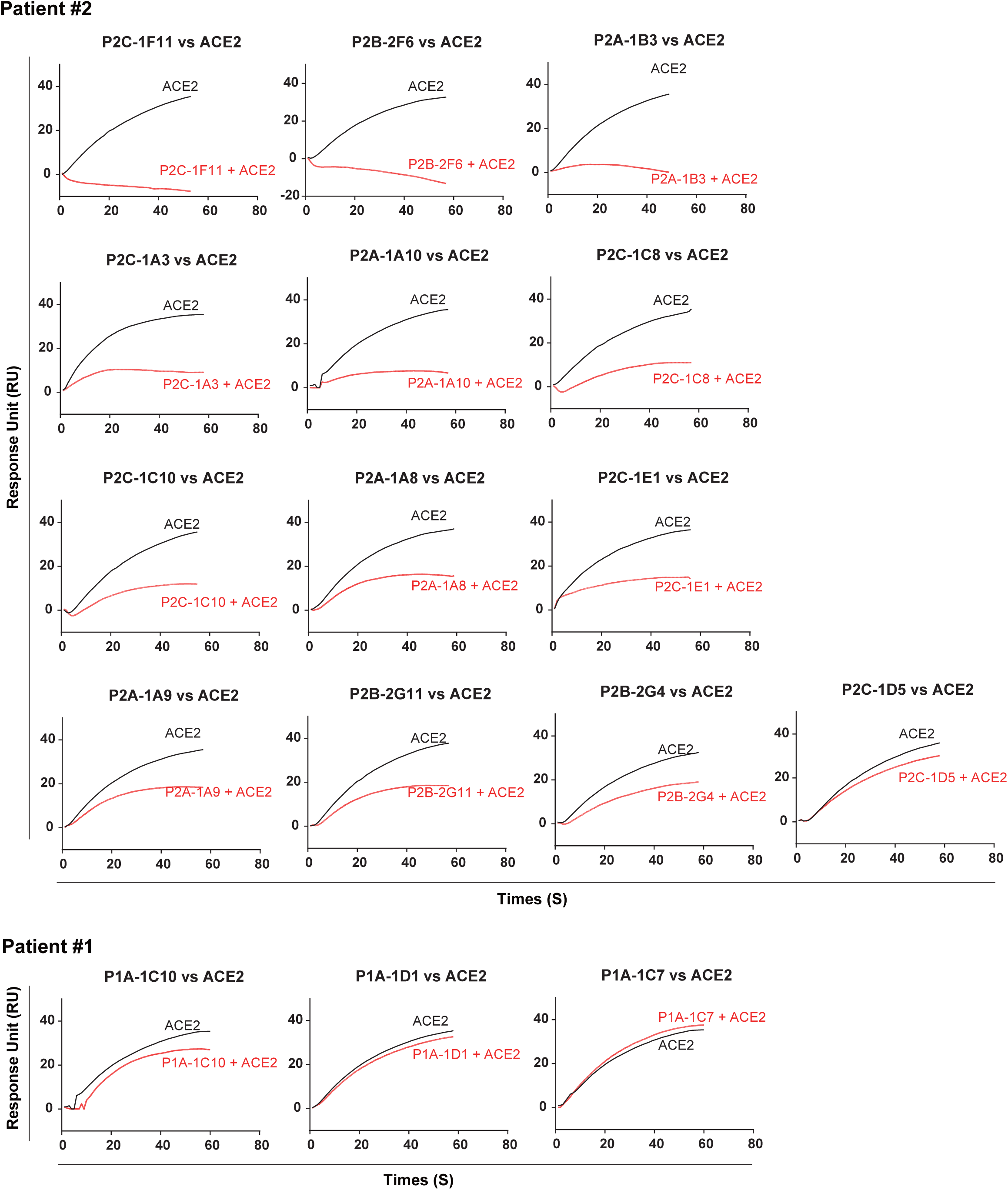
Antibody and ACE2 competition for binding to SARS-CoV-2 RBD measured by SPR. The sensorgrams show distinct binding patterns of ACE2 to SARS-CoV-2 RBD with (red curve) or without (black curve) prior incubation with each testing antibody. The competition capacity of each antibody is indicated by the level of reduction in response unit of ACE2 comparing with or without prior antibody incubation.

**Figure S6.**
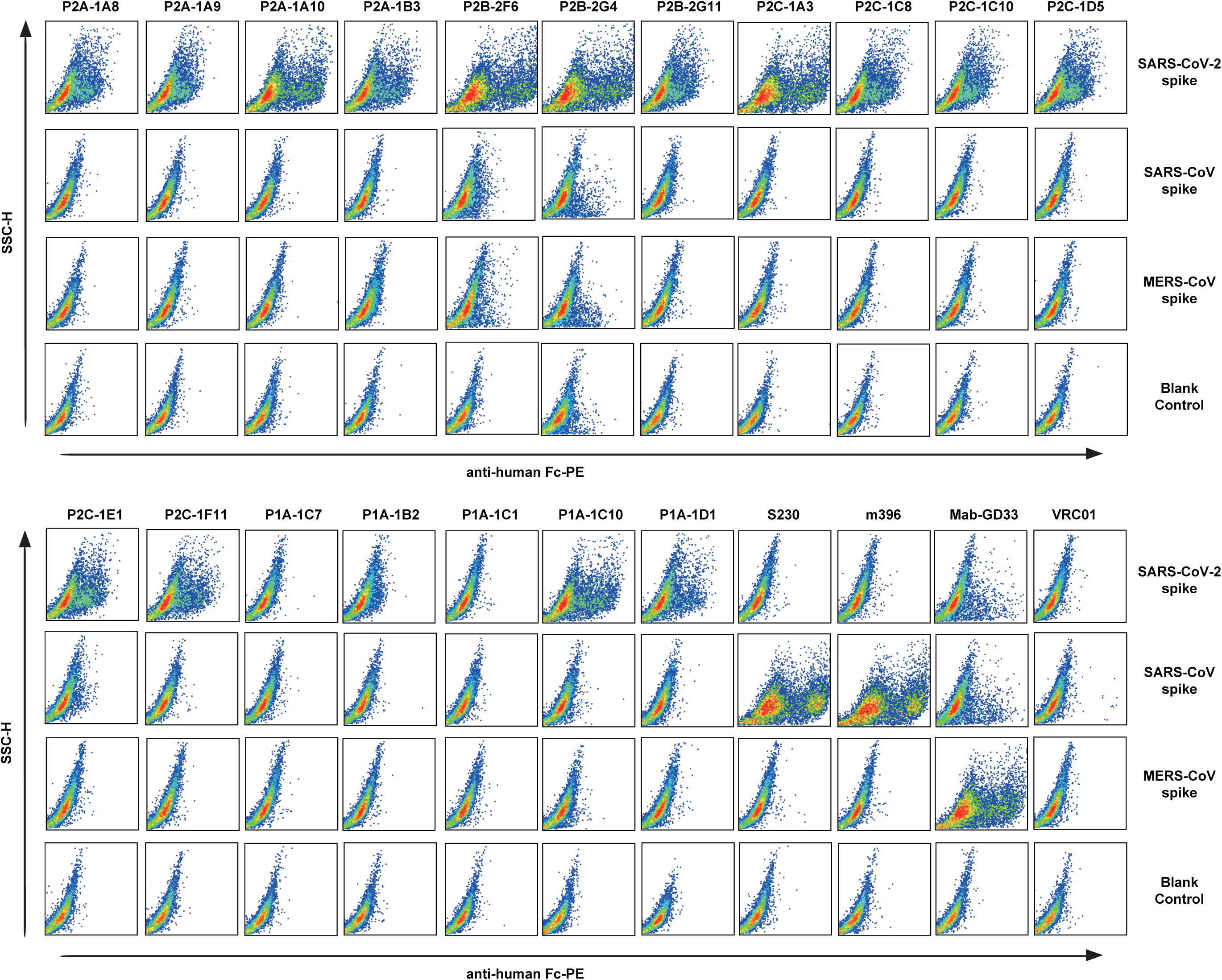
Analysis of antibody binding to cell surface expressed trimeric Spike protein. HEK 293T cells transfected with expression plasmid encoding the full length spike of SARS-CoV-2, SARS-CoV or MERS-CoV were incubated with 20ug/ml testing antibodies. The cells were then stained with PE labeled anti-human IgG Fc secondary antibody and analyzed by FACS. Positive control antibodies include S230 and m396 targeting the RBD of SARS-CoV Spike, and Mab-GD33 targeting the RBD of MERS-CoV Spike. VRC01 is the negative control antibody targeting HIV-1 envelope glycoprotein.

**Figure S7.**
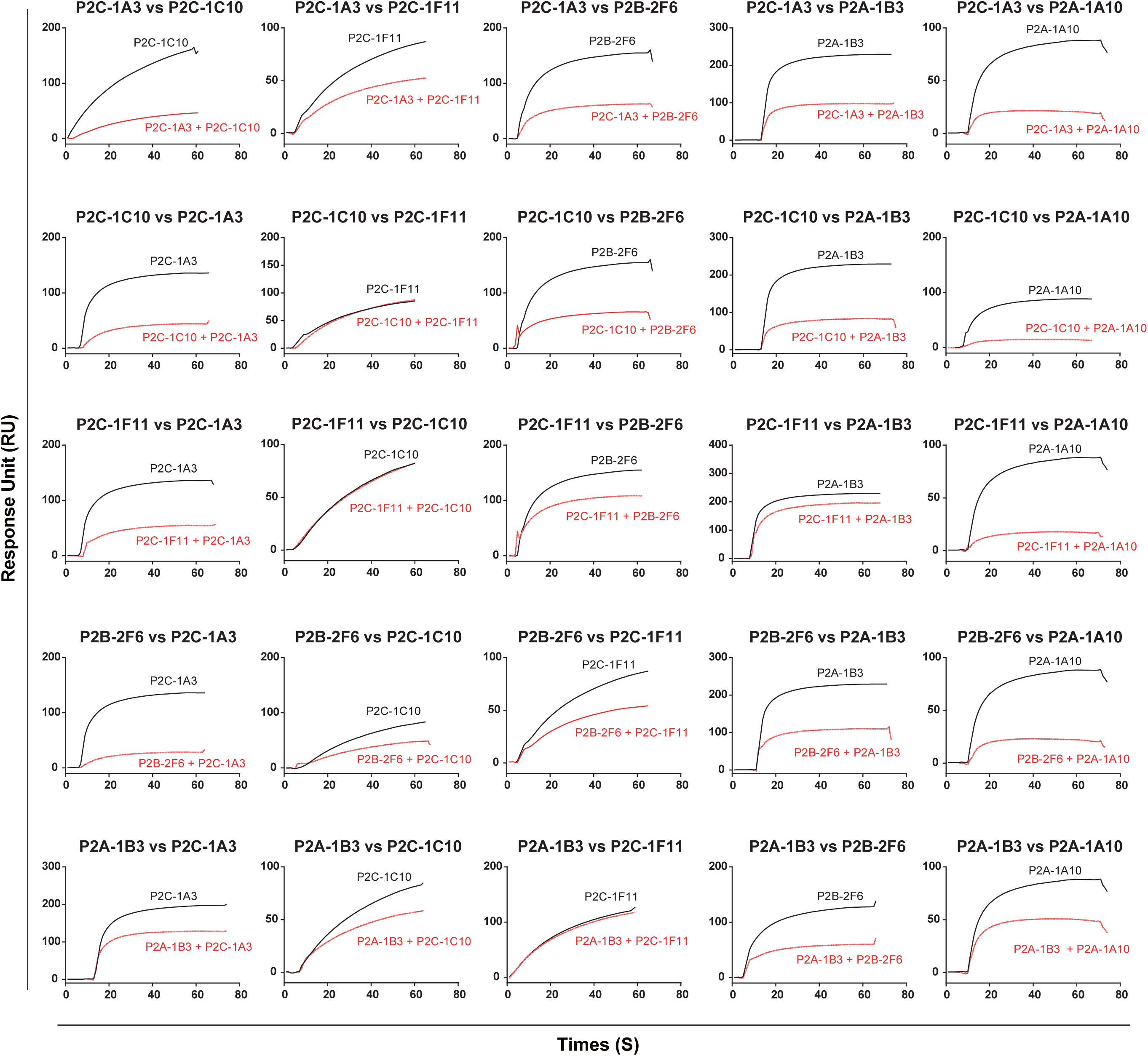
Epitope mapping through competitive binding measured by SPR. The sensorgrams show distinct binding patterns when pairs of testing antibodies were sequentially applied to the purified SARS-CoV-2 RBD covalently immobilized onto a CM5 sensor chip. The level of reduction in response unit comparing with or without prior antibody incubation is the key criteria for determining the two mAbs recognize the separate or closely situated epitopes.

**Table S1.**
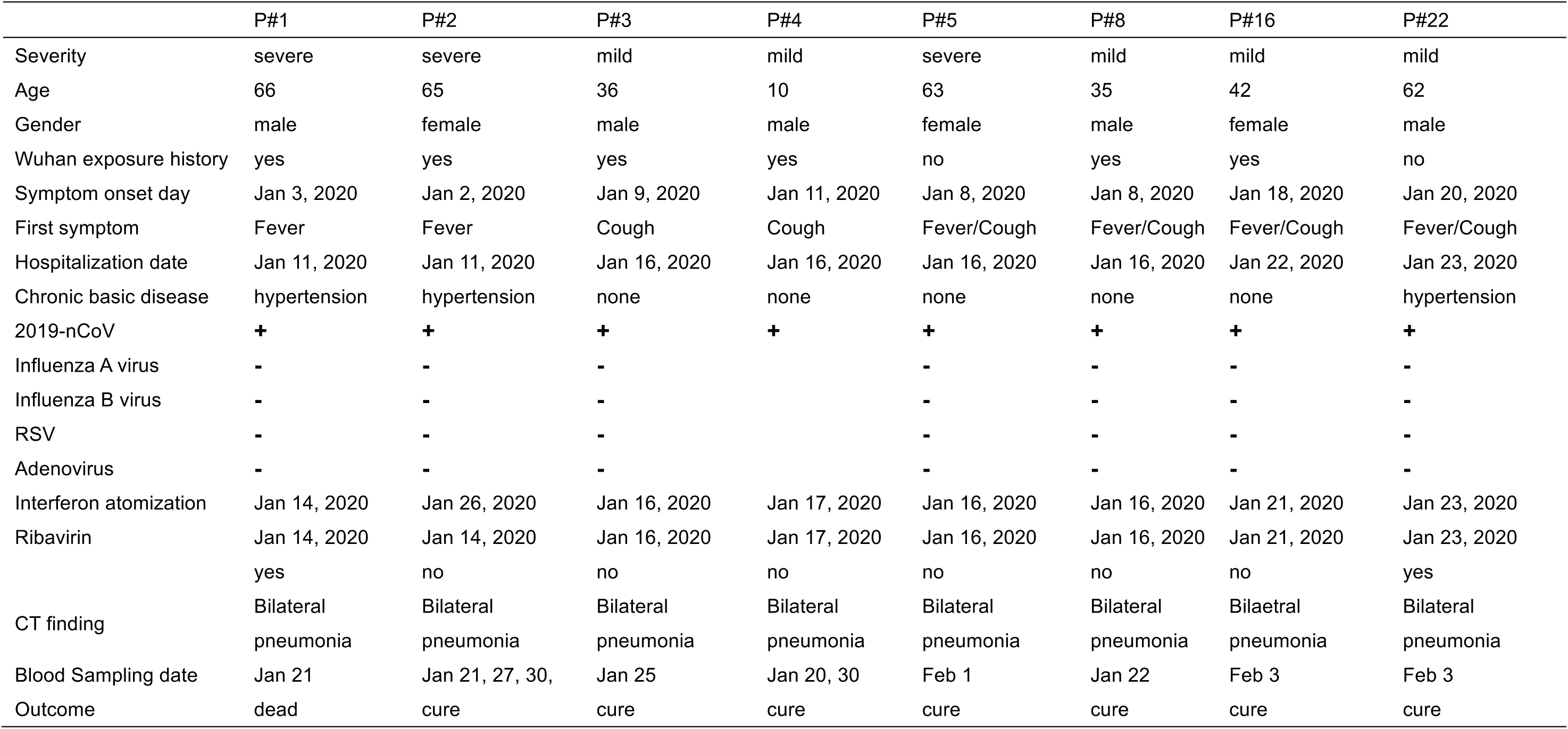
Clinical characterization of the study subjects.

**Table S2.**
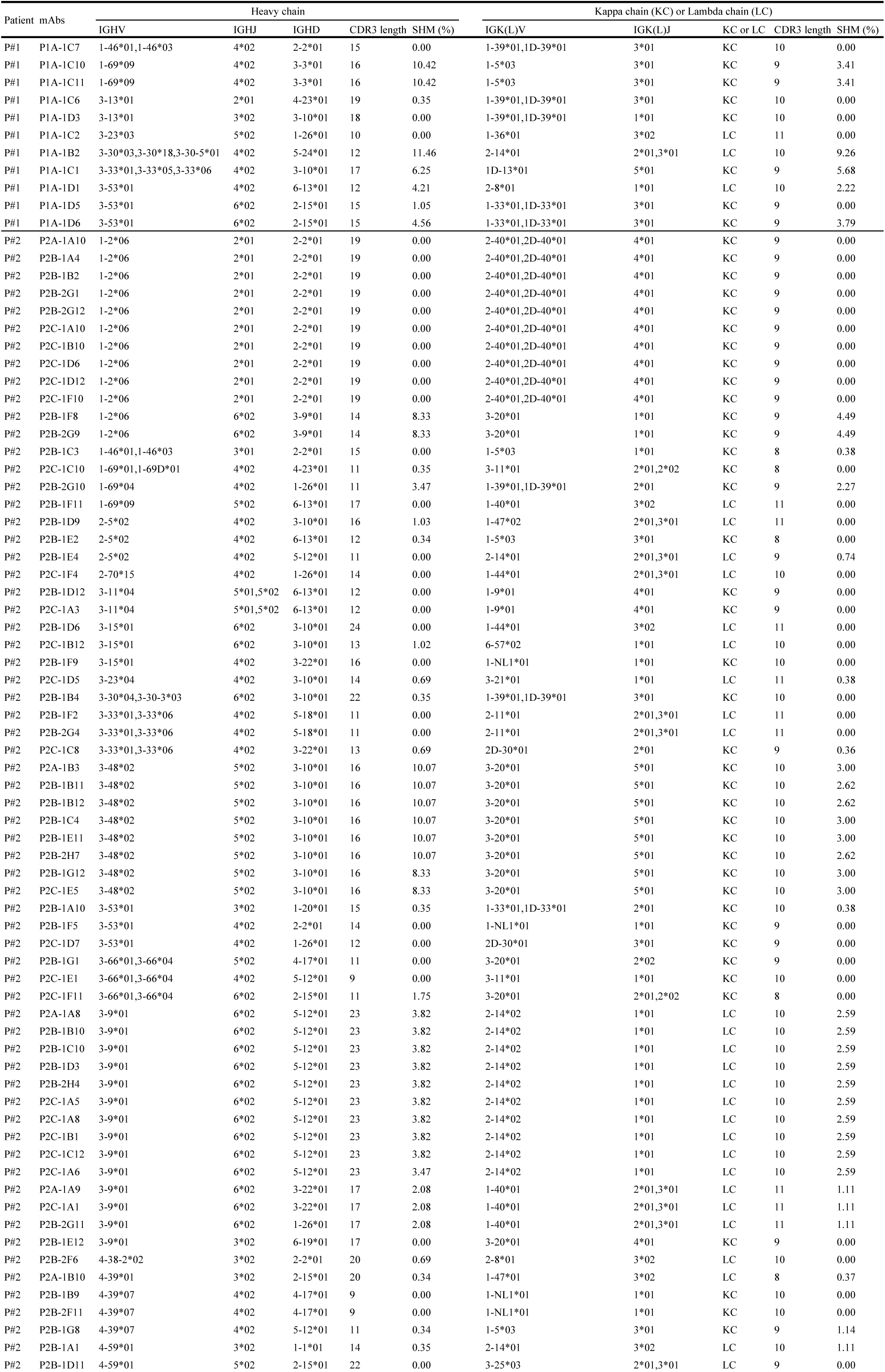
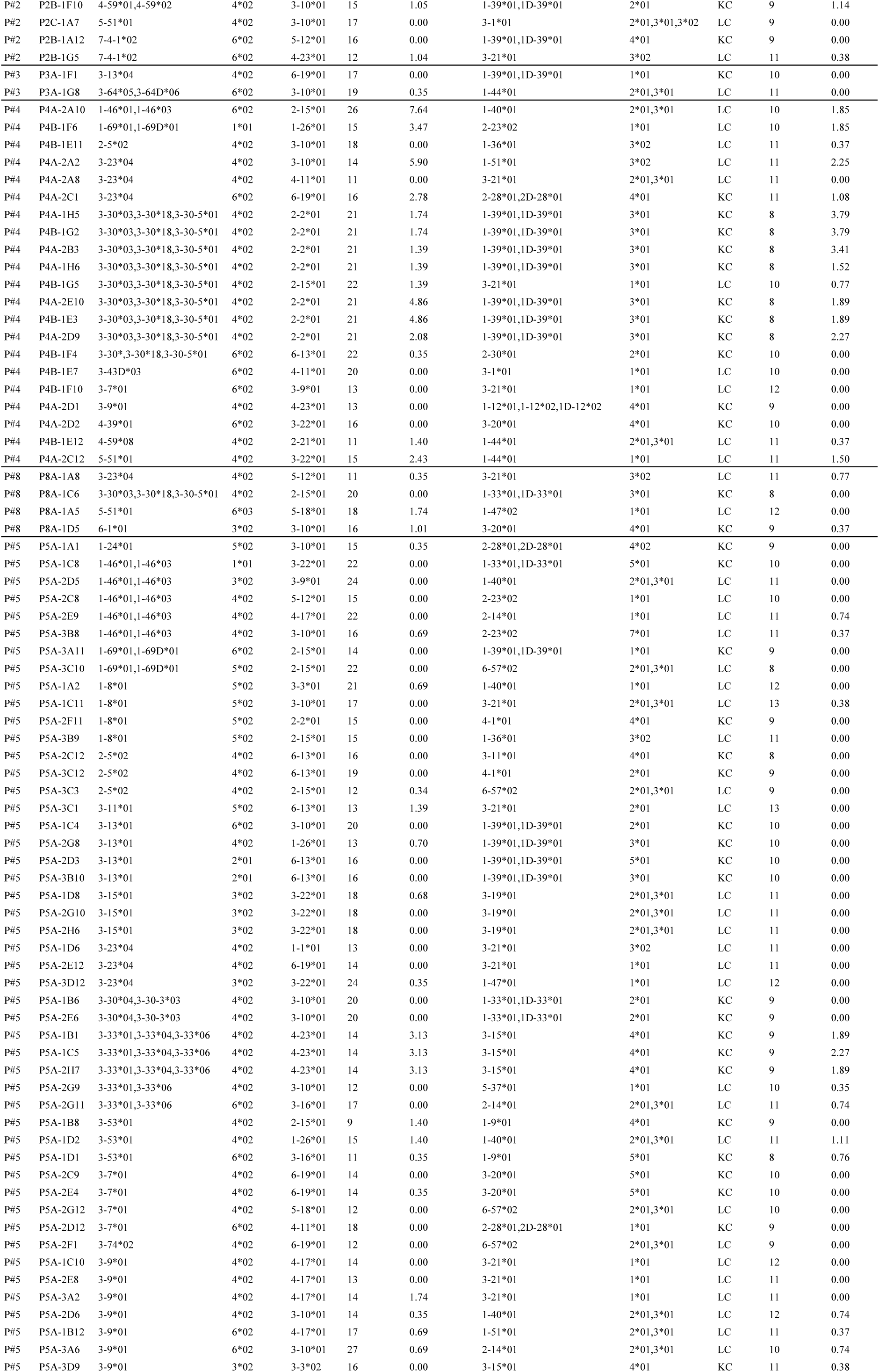
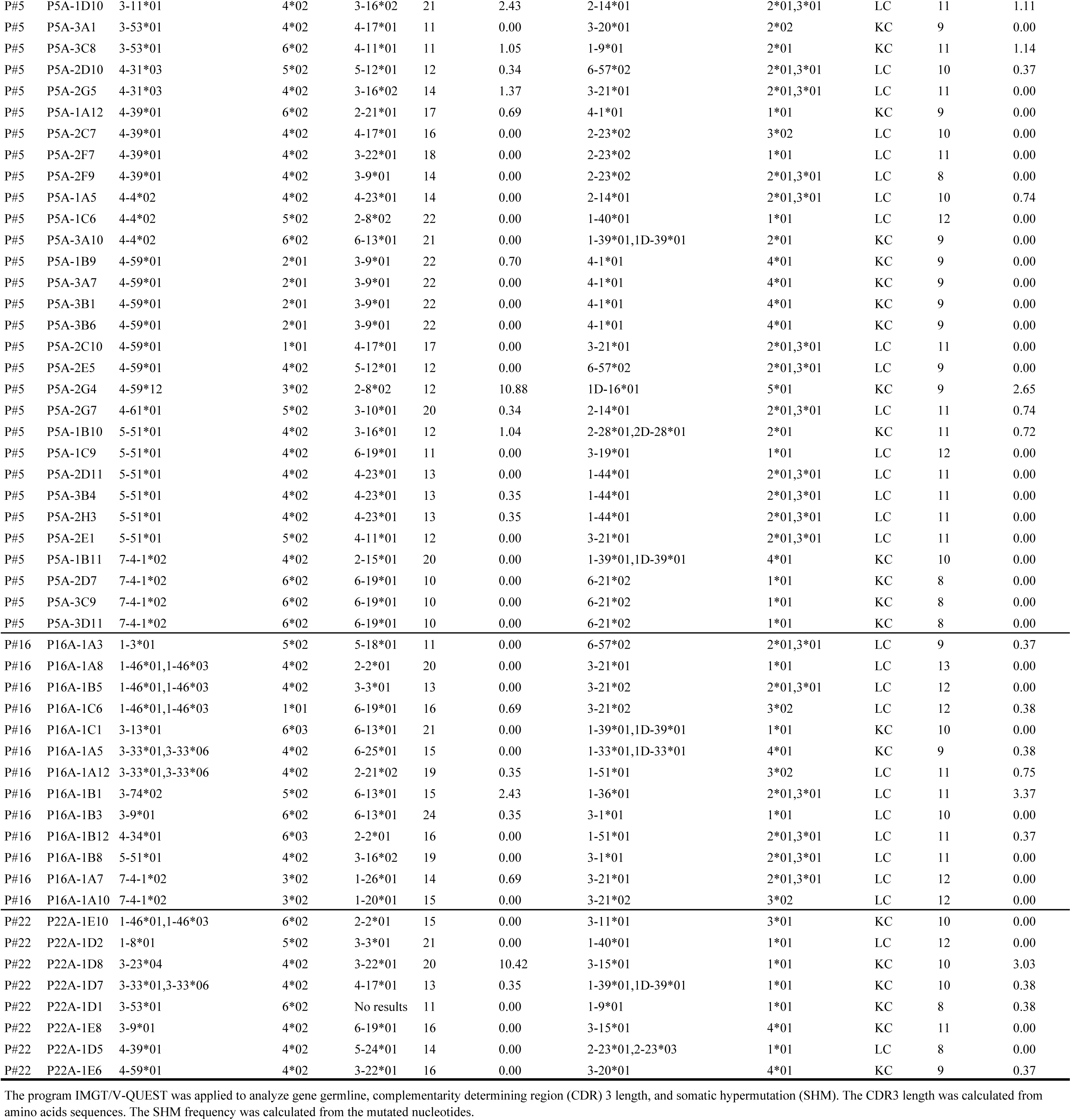
Gene family analysis of monoclonal antibodies.

